# Telomerase modRNA offers a novel RNA-based approach to treat human pulmonary fibrosis

**DOI:** 10.1101/2025.02.19.639041

**Authors:** Jia Li Ye, Klaudia Grieger, Dongchao Lu, Christina Brandenberger, Malte Juchem, Maria Jordan, Lea Oehlsen, Patrick Zardo, Christopher Werlein, Christina Hesse, Katherina Sewald, Sandy Tretbar, Thomas Thum, Shambhabi Chatterjee, Christian Bär

## Abstract

Pulmonary Fibrosis (PF) is a life-threatening illness that is characterized by progressive scarring in the lung interstitium. There is an urgent need for new PF therapies because current treatments only slow down the progression of fibrosis and the median life expectancy post-diagnosis is only 4 to 6 years. Since PF patients frequently exhibit telomere attrition, overexpressing telomerase, the enzyme responsible for synthesizing telomeres represents a compelling therapeutic option. In this study, we *in vitro* transcribed human telomerase reverse transcriptase (hTERT) mRNA using modified nucleosides (modRNA). ModRNA hTERT treatment led to transient activation of telomerase activity in a dose-dependent manner in MRC-5 cells and, importantly, in primary human alveolar type II pneumocytes (ATII cells). Consequently, the proliferative capacity was increased, concomitant with reduced DNA damage and elongated telomere length. Notably, induction of cellular immune response was only detectable at the highest modRNA concentration, and returned to normal levels within 48 hours. Next, we demonstrated that circularized, exonuclease-resistant modRNA hTERT extended the transient expression profile which may be clinically advantageous. Finally, we provided therapeutic proof of concept in organotypic 3D *ex vivo* human precision-cut lung slices derived from end-stage PF patients. Intriguingly, a single modRNA hTERT treatment inhibited senescence as indicated by significantly lower levels of senescence-associated β-galactosidase, and pro-inflammatory IL6 and IL8. Concurrently, the key fibrosis mediators TGFβ and COL1A1 were markedly reduced.

In conclusion, the data presented herein provide initial evidence for the potential of RNA-based hTERT therapy for treating human lung fibrosis.

## Introduction

Lung diseases arise due to a variety of factors such as viral or bacterial infection, asthma, bronchitis, fibrosis, air pollution or occupational risks affecting specific cell types of the lung. Interstitial lung diseases (ILD) comprises of several heterogeneous conditions which ultimately lead to inflammation and scarring of the lung tissue. Out of these pulmonary fibrosis (PF) is a chronic fatal sub-type of ILD, marked by remodeling of the alveolar compartment, progressive fibrosis in the interstitium, and hindrance of a proper gas exchange, ultimately resulting in respiratory failure (Cottin et al., 2018; Fan, Gui, Zhou, Li, & Chen, 2024). In addition, a worse survival prognosis of only 4 to 6 years post-diagnosis (Cottin et al., 2018; Khor et al., 2023; Strand et al., 2014; Yamazaki, Nishiyama, Yoshikawa, Tohda, & Matsumoto, 2022), put these lung patients at high risk. There are currently two FDA-approved drugs, pirfenidone and nintedanib which show potential to slow down the fibrotic progression but fail to reverse or cure the disease, highlighting an urgent need to discover novel therapeutic strategies against PF (Cilli & Uzer, 2023; Wijsenbeek & Cottin, 2020).

Telomere dysfunction, a hallmark of ageing, has also been recognized as a major risk factor for both PF and its sub-type idiopathic pulmonary fibrosis (IPF) pathology caused by chronic alveolar epithelial injury, senescence, and tissue remodeling (Alder & Armanios, 2022; McDonough et al., 2018; Newton et al., 2016; Piñeiro-Hermida et al., 2022; Van Batenburg et al., 2020). The ageing population is most affected by IPF, with an overall prevalence of 10-60 cases per 100,000 individuals, which drastically increased to 400 cases per 100,000 in individuals over 65 years of age (Esposito et al., 2015; Raghu et al., 2014). There is a high correlation between telomere length and IPF disease severity (Adegunsoye et al., 2023; Duckworth et al., 2021). Around 25% of individuals with familial pulmonary fibrosis (FPF) exhibit mutations in telomere maintenance genes, such as TERT, TERC, DKC1, TIN2 and RTEL1, predisposing these patients to senescence and subsequently early alveolar epithelial cell failure (Peljto et al., 2023; Podolanczuk et al., 2023). Notably, those with sporadic IPF without any known mutations in the telomere machinery, possess elevated levels of telomere shortening too; indicating that senescence induced by shortened telomeres is a unifying factor of both the sporadic and FPF forms of IPF (Cronkhite et al., 2008). Mechanistically, telomere shortening and dysfunction in alveolar type II pneumocytes (ATII cells) have been shown to promote DNA damage and cellular senescence (Alder et al., 2015; Van Batenburg et al., 2021; Yazicioglu et al., 2020). In addition to producing surfactant, the ATII cells also serve as progenitors for alveolar type I pneumocytes (ATI cells). Hence, ATII cell senescence results in limited regeneration capacity after lung injury (Hirsch et al., 2024; Povedano, Martinez, Flores, Mulero, & Blasco, 2015). Moreover, senescent ATII cells secrete pro-fibrotic factors which promote the development of PF. Ultimately, the progressive expansion of fibroblast and myofibroblasts leads to extracellular matrix remodeling (Spagnolo et al., 2021).

Considering that telomere shortening is a key driver and strongly associated with PF, reactivating the enzyme telomerase which synthesizes telomere repeats would evidently offer a powerful therapeutic approach to replenish telomeres, and to mitigate cellular senescence in PF (Povedano et al., 2018; Bär & Blasco, 2016). Promising results after reactivation of telomerase reverse transcriptase (TERT), catalytic subunit of telomerase, have been shown in a mouse model of bleomycin induced PF. AAV9-Tert-mediated gene therapy showed improvement in lung volume and mechanics as well as lower inflammation concomitant with improved regression of the fibrosis (Povedano et al., 2018). Despite successful TERT reactivation using gene therapy tools in mice models, AAV vectors present some concerns regarding clinical application due to the slow kinetics of transgene expression and the long-term and constitutive TERT overexpression which imposes risk of tumorigenesis (Penaud-Budloo et al., 2008). Alternatively, the use of an mRNA encoding human telomerase reverse transcriptase (hTERT) enables a much faster and transient reactivation of telomerase activity which might be sufficient to achieve telomere elongation, thus offering a safer and efficient therapeutic approach. If needed, the transient RNA-mediated hTERT reactivation may be extended by utilizing circularized versions of the mRNA or self-amplifiying mRNAs which offer higher stability and longer expression (Wesselhoeft, Kowalski, & Anderson, 2018).

In this study, we utilized modRNA hTERT to reactivate telomerase activity in primary human ATII cells, which led to improved cellular fitness and telomere elongation. We demonstrated that circularized RNA hTERT is functional and extends the window of transient telomerase activity. Finally, we utilized precision-cut lung slices (PCLS) from end-stage lung fibrosis patients to provide the first proof-of-principle for the anti-senescent and anti-fibrotic effectiveness of hTERT modRNA therapy in human lung fibrosis.

## Methods

### Human lung material

Human primary ATII cells (EP71AL Epithelix) were obtained from non-smoking donors with no reported pathology (Epithelix Sarl, Geneva, Switzerland).

Human Precision-cut lung slices (PCLS) of end-stage PF patients were diagnosed with usual interstitial pneumonia (UIP) pattern and were provided by the Hanover Medical School (MHH, Hanover, Germany). Patients’ demographic and diagnosis are summarized in Table 1. The experiments with human lung tissue were approved by the ethics committee of the Hannover Medical School (Hannover, Germany) in compliance with “The Code of Ethics of the World Medical Association” (renewed on 2015/04/22, number 2701–2015). All patients or their next of kin, gave written informed consent for using their lung tissue for research.

**Table 1:** Patients’ demographic and diagnosis.

| Gender & Age [years] | Diagnosis | Corresponding Figure(s) |
| --- | --- | --- |
| Female, 64 | Tumor | Fig. 4B, Supplement Fig. 4A |
| Female, 58 | UIP, EAA fibrosis | Fig. 4B, Supplement Fig. 4A |
| Male, 49 | UIP; secondary fibrosis | Fig. 4C-F |
| Male, 53 | UIP; secondary fibrosis | Fig. 4B-F |
| Male, 61 | UIP; IPF | Fig. 4C-F |
| Male, 56 | UIP; IPF | Fig. 4C-F |

UIP: usual interstitial pneumonia; EAA: Extrinsic allergic alveolitis.

### Cell culture, quantitative growth curve and transfection

Human primary fetal lung cells (MRC-5 cell line) and HEK293 cells were cultured in high glucose DMEM (41965062 GibcoTM) with 10 % fetal bovine serum and 1 % penicillin/ streptomycin. Human primary ATII cells were cultured with ready-to-use Epithelix Culture Media. Cells were counted with the Countess^TM^ II from Invitrogen. To generate the growth curve, population doubling level (PDL) was calculated by using the equation PDL = 3.32 · (log n2 - log n1) + x, where n1 and n2 are the respective population size and x is the PDL from the previous passage. For transfection, Lipofectamine RNAiMAX (133778075 Thermo Fisher Scientific) was mixed with 500 ng modRNA in Opti-MEM in a 2:1 vol:vol ratio and added to the cells in a 0.5 mL transfection volume to achieve a final treatment concentration of 1 µg/mL treatment. For dose dependent r concentration, the amounts were adjusted accordingly. After 4 h culture media (1.5x of from the total transfection volume) was added onto the transfected cells.

### Precision-cut lung slices

Precision-cut lung slices were generated based on previously described protocols (Hesse et al., 2022; Switalla et al., 2010). After an overnight recovery period, PCLS were treated with Lipofectamine RNAiMAX (133778075 Thermo Fisher Scientific) and 500 ng modRNA in Opti-MEM in a 2:1 vol:vol ratio for 4 h. Afterwards 1.5x culture media (of the total transfection volume) was added onto the transfected cells for subsequent culture at 37 °C and 5 % CO_2,_ with a medium exchange 48 h post-transfection. Culture media for PCLS comprised of DMEM/F12 (11039021 Gibco) and 1 % penicillin/ streptomycin.

### Lactate Dehydrogenase (LDH) assay

Tissue viability was assessed using the lactate dehydrogenase (LDH) based Cytotoxicity Detection Kit (11644793001 Roche) according to the manufacturer’s recommendations.

### Enzyme-linked immunosorbent assay (ELISA) assay

PCLS culture supernatants were collected after 96 h, supplemented with 0.2 % protease inhibitor cocktail (87785 Thermo Fisher Scientific), and stored at −80 °C. Human IL6, IL8, TGFβ and pro-COL1A1 was measured using DuoSet ELISA kits from R&D Systems (DY206, DY208, DY6220 and DY240) according to the manufacturer’s recommendations.

### Senescence-associated β-galactosidase staining

Senescence-associated β-galactosidase (SA-β-gal) levels were assessed within the tissue after 96 h using a β-Gal Staining Kit (9860 Cell Signalling) and performed according to the manufacturer’s recommendations adjusting staining/fixation volumes to 500 µL per PCLS. Overview Images of the stained PCLS were recorded using a stereomicroscope (Discovery V8; Zeiss, Germany) controlled by the Axio Vision 4.8.2 software program (Zeiss, Germany). Afterwards the tissue was embedded in OCT (6478.2 Roth), frozen to -20 °C and sectioned into 10 µm-thick slices. Cryo sections were fixed on SuperFrost microscope slides (J1800AMNZ Epredia) with DPX mounting medium (10021203 Sigma Aldrich). Embedded β-gal stained PCLS slides were imaged with a 20X objective (PlanApo 20x 0.75/0.60 mm) at a fluorescence microscope (BZ-X810 Keyence). Images were stitched together and quantified using the BZ-X800 Analyzer Software.

### mRNA template design, synthesis and validation

#### Linear construct

To generate the mRNA construct, the open reading frame (ORF) of the respective gene (hTERT, GFP) was cloned into the multiple cloning site of a pMA-RQ plasmid containing a CleanCap®-adjusted T7 promotor (TAATACGACTCACTATAAG), 5′ untranslated region (UTR) human α-globin, 3′ UTR human β-globin, segmented poly-A tail (60A-G-60A) and a BspQI linearization site. Before modRNA production, the plasmid was sequenced, linearized and then the *in vitro* transcription (IVT) was performed to produce modRNA. For IVT productions a final concentration of 5 mM for each rATP, rCTP, rGTP and m1Ψ, 4 mM CleanCap AG (N-7113-1 TriLink BioTechnologies), transcription buffer containing 40 mM Tris-HCl (pH 8), 10 mM dithiothreitol (DTT), 2 mM spermidine, 0.002 % Triton X-100, 16.5 mM magnesium acetate, 1 U/µL RNA inhibitor, 0.002 U/µL inorganic pyrophosphatase and 8 U/µL T7 RNA polymerase was used. After 4 h incubation at 37 °C, the IVT product was treated with 0.04 U/µL TURBO^TM^ DNase (AM2239 Thermo Fisher Scientific) for 15 min at 37 °C, filtered with Amicon® Ultra-4 (UFC801024 Merck), 5 U/µL Antartic Phosphatase (M0289L New England Biolabs) for 1 h at 37 °C treated and purified again with Monarch® RNA Cleanup Kit (T2050L New England Biolabs). The final modRNA product was verified with a 1 % denaturing agarose gel electrophoresis.

#### Circular construct

The hTERT ORF was cloned into the multiple cloning site, containing T7 promotor, group I self-splicing intron and IRES (internal ribosomal entry site) sequence. Splicing strategy was adapted as previously described (Wesselhoeft et al., 2018). To generate circular RNA, 400 ng of the sequenced plasmid was used for a PCR using a master mix consisting of 1X Phusion® High Fidelity PCR Master Mix with GC Buffer (M0532 New England Biolabs), 1 µM primer pair (fw: TAATACGACTCACTATAGGG GGAGA; rev: GTTTAAACGGGCCCTCTAGACTCGAG) and 10 % DMSO.

Thermocylcer protocol included the following steps of 98 °C for 30 s; 30 cycles of 98 °C for 5 s, 64 °C for 10 s, 72 °C for 2 min; 72 °C for 5 min. After purification with QIAquick PCR Purification Kit (28106 QIAGEN) the IVT protocol as described above was performed with an overnight (12-16 h) incubation. CleanCap AG was not used and m1Ψ was substituted by rUTP. Purification was performed, followed by 20 U/µL RNase R (172010 Biozym) treatment for 30 min at 37 °C and repeated purification with Monarch® RNA Cleanup Kit (T2050L New England Biolabs). The final product was verified with sequencing, 1 % denaturing agarose gel electrophoresis and additionally reverse transcription followed by PCR with divergent primer pairs (primer list provided in Table 2).

**Table 2:**
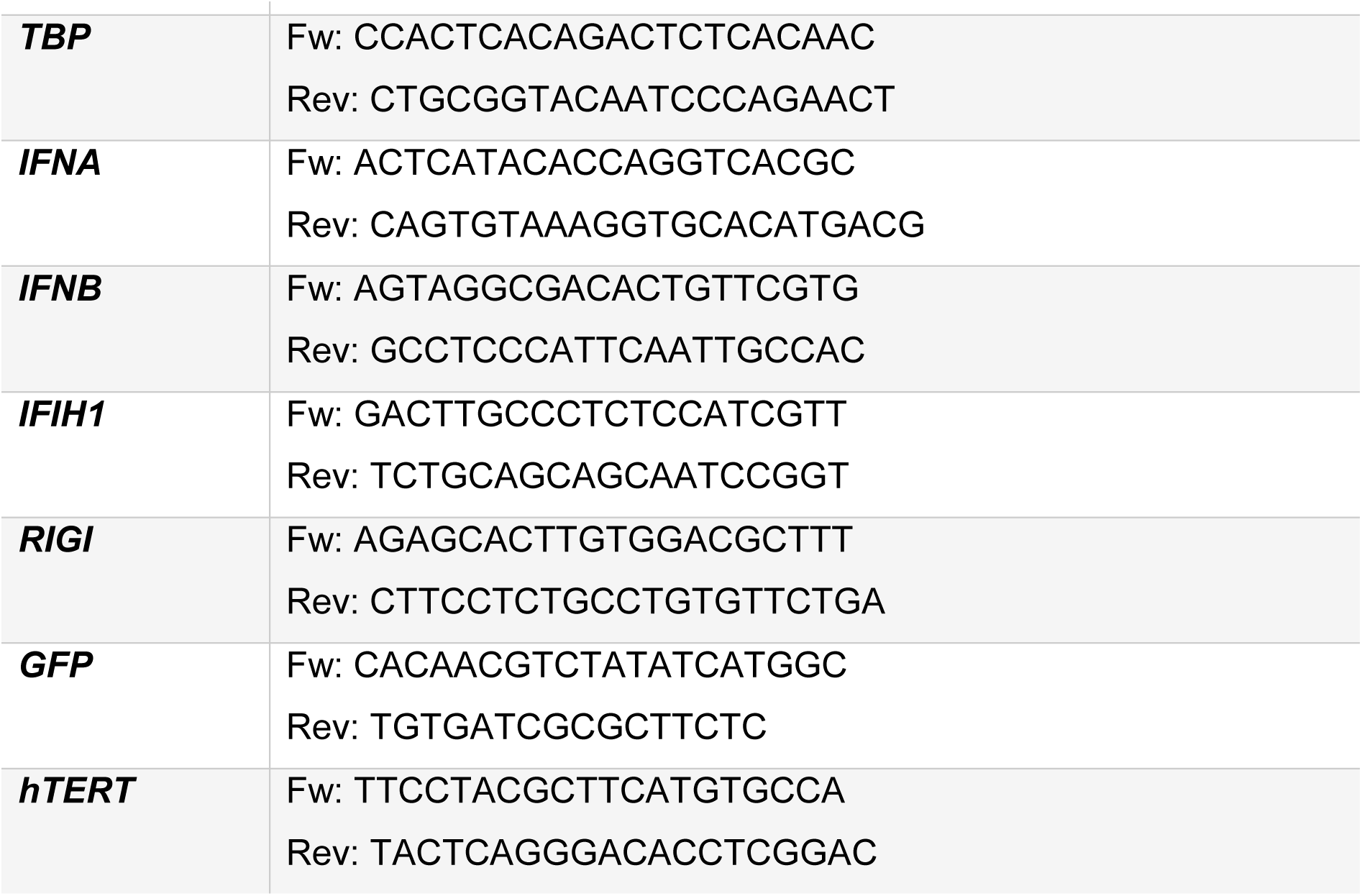
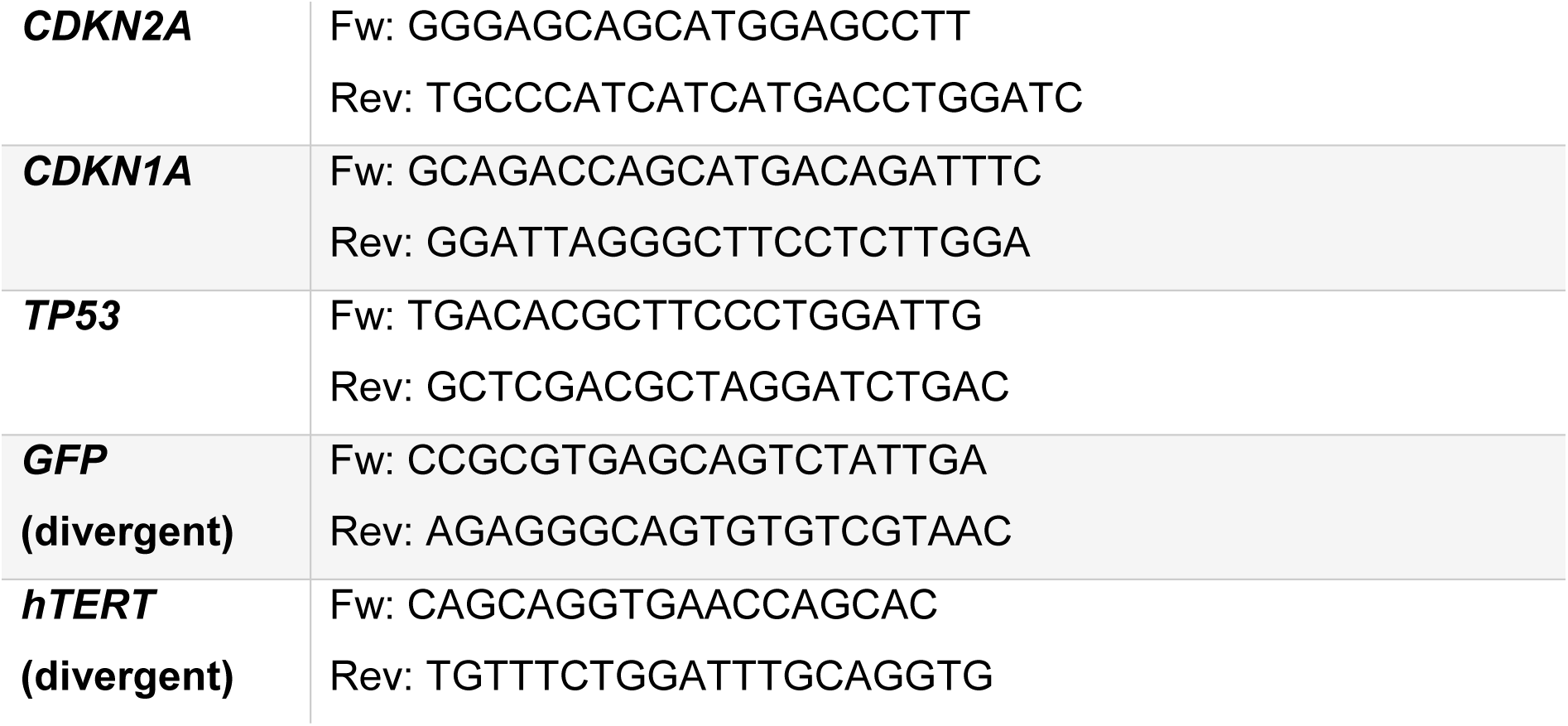
List of genes and their corresponding primers in the qPCR and PCR.

### RNA isolation and quantitative PCR

PCLS RNA isolation was performed based on previously described protocol (Niehof et al., 2017). For all the other samples in this study, RNA was isolated using QIAzol Lysis Reagent (79306, QIAGEN), precipitated in 75 % ethanol and eluted in DNase/RNase free H_2_O. The RNA concentration was measured with Take3 Plates on a Bio-Tek plate reader (Synergy HT). To remove all DNA contamination, 500 – 1000 ng RNA was treated with 0.2 U/µL RNase-Free DNase (79254 QIAGEN) for 30 min at 37 °C, followed by reverse transcription by using Biozym cDNA synthesis kit (331470 Biozym). The real-time PCR was performed on QuantStudio^TM^ 7 Flex System (Thermo Fisher Scientific) with specific primer pairs (Table 2).

### BrdU incorporation assay

Cell Proliferation ELISA, BrdU (11647229001, Merck) was used according to the manufacturer’s instructions.

### Immunostaining

Cultured cells were shortly washed with PBS and fixed with 4 % paraformaldehyde (PFA) by incubating for 10 min, followed by three times washing with PBS. Afterwards cells were permeabilized with 0.1 % Triton X-100 for 10 min, followed by three times washing with PBS. 5 % donkey serum in PBS (blocking buffer) was added for 30 min followed by the primary antibody (rabbit-anti-Ki67 1:250 ab16667 abcam, mouse-anti-pH3 1:400 #9706 Cell Signaling Technology, rabbit-anti-γH2AX 1:2,000 ab11174 abcam). Primary antibodies were diluted in blocking buffer and incubated at 4 °C overnight. Cells were washed three times with PBS and then incubated with the 1:1,000 diluted secondary antibody (donkey-anti-rabbit Alexa Fluor^TM^ 488 A21206 Invitrogen, donkey-anti-mouse Alexa Fluor^TM^ 488 A21202 Invitrogen) and 1:1,000 diluted Hoechst for 30 min. Next, the cells were washed three times with PBS. Images were acquired with Cytation 1 (Biotek) and analyzed with the BioTek Gen5 software.

### Telomerase repeat amplification protocol (TRAP)

The protocol was adapted from a previously described protocol (Herbert, Hochreiter, Wright, & Shay, 2006). 200,000 cells (based on live cell count) were harvested, washed with PBS and centrifuged at 5,000 g for 5 min at 4 °C. Then the supernatant was removed, the cell pellet was snap frozen in liquid nitrogen and stored at -70 °C until further processing. To lyse the cells ice-cold NP-40 lysis buffer was added to a final concentration of 1,000 cells/µL and incubated for 30 min on ice. 2 µL cell lysate was added to 48 µL master mix. The master mix contained 50X primer Mix (100 ng/mL each of ACX and NT primers along with 0.001 attomol/mL TSNT primers), DY-682 labelled TS primer (Table 3). Additionally, each sample had a respective heat-inactivated (10 min at 85 °C) cell lysate as a quality control for the TRAP assay. The TRAP PCR was run on a thermocycler with following protocol: 25 °C for 30 min; 95 °C for 5 min; 24 cycles of 95 °C for 30 s, 52 °C for 30 s, 72 °C for 40 s . Loading buffer was added to each sample before loading on a 10 % acrylamide gel (19:1 Acryl: Bis acrylamide in Tris-borate-EDTA buffer) and the gel was run for around 3.5 h at 250 V. The gel was fixed for 15 min in fixative solution (0.5 M NaCl, 50 % EtOH, 40 mM Sodium Acetate at pH 4.2) and then transferred in H_2_O right before scanning with LICOR Odyssey Imaging System 9120.

**Table 3:**
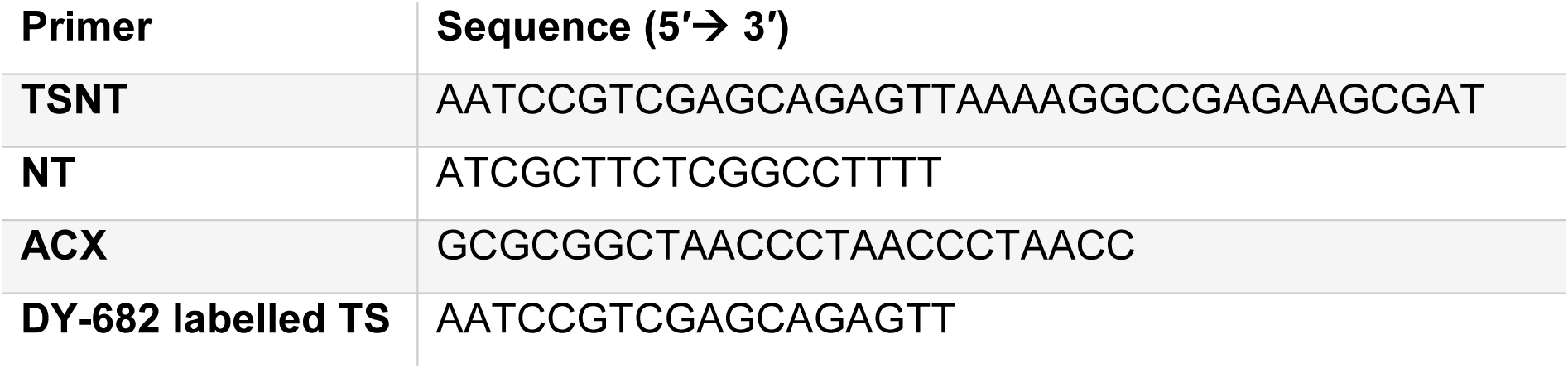
List of primers used in TRAP assay.

### Quantitative fluorescence *in situ* hybridization (qFISH)

The protocol was adapted from a previously described protocol (O’Sullivan et al., 2005). First cultured cells were washed with PBS and then 150,000-300,000 cells were spotted onto SuperFrost microscope slides (J1800AMNZ Epredia) using a Thermo Shandon Cytospin 3 centrifuge. The slides were stored at 4 °C until all samples were collected for staining. For the staining, the slides were washed twice with PBS for 15 min and then fixed in 4 % PFA in PBS for 2 min. After washing the slides thrice with PBS for 5 min, slides were incubated with pre-warmed pepsin in a 37 °C water bath for 10 min. Followed by washing twice in PBS and again fixed in 4 % PFA for 2 min. Afterwards slides were washed thrice in PBS for 5 min. To dehydrate the cells, slides were incubated in different EtOH concentrations (70 %, 90 % and 100 %) for 5 min each continued by air-drying until no drops were visible. 15-30 µl of Telomerase PNA probe mix (10 mM TrisCl pH 7, 25 mM MgCl_2_, 9 mM citric acid, 82 mM Na_2_HPO_4_, 7 % deionized formamide, 0.25 % blocking reagent and 0.5 mg/mL Telomeric PNA probe) were spotted onto the slides and sealed with cover slips. These slides were incubated at 85 °C for exactly 3 min. Subsequently, the slides were incubated in a dark and wet chamber for around 4 h. Next, the cover slip was removed and washed twice for 15 min each in 10 mM TrisCl pH 7.2, 0.1 % BSA in H_2_O and 70 % formamide under vigorous shaking (550 rpm). Next, slides were washed thrice for 5 min with TBS-T and once for 5 min with PBS. The slides were incubated for 20 min with 1:1,000 Hoechst in the dark and then washed thrice for 5 min with PBS. In the end, the slides were air-dried and around 10 µL Prolong^TM^ Gold Antifade Mountant (P10144 Thermo Fisher Scientific) was dropped onto the slides and sealed with cover slips. The slides were stored in the dark at 4 °C. The images were acquired using 63X oil objective and additional 2X zoom with 405 nm and 561 nm laser in the confocal microscope Zeiss LSM 780. Image analysis was performed using our own software programmed for quantification of the digitized fluorescent signals. The software is based on the principals of the open source Telometer Plugin. The software was developed in C# with the integrated development environment Visual Studio Community Edition 2019 under Microsoft .NET and supplemented with OpenCV routines via the EmguCV.NET wrapper (EmguCV 2023).

### Transmission electron microscopy

Cell culture inserts (10482885 Thermo Fisher Scientific) with ATII cells were treated with Lipofectamine RNAiMAX (133778075 Thermo Fisher Scientific) and 500 ng modRNA in Opti-MEM in a 2:1 vol:vol ratio. After 4 h culture media (1.5x of the total transfection volume) was added onto the transfected cells. Cells were fixed using a mixture of 1.5 % paraformaldehyde and 1.5 % glutaraldehyde in 0.15 M HEPES buffer for at least 24 h. The samples were then embedded in epoxy resin as described previously (Brandenberger, Kling, Vital, & Mühlfeld, 2018; Kling, Lopez-Rodriguez, Pfarrer, Mühlfeld, & Brandenberger, 2017) and cut into ultrathin section for transmission electron microscopic (TEM) analysis. Imaging was done with Morgagni 268 microscope (FEI).

### Statistics

Batch correction was performed for all experiments, besides growth curve, immunostaining, qFISH and SA-β-gal analysis before statistical analysis. All data were analyzed with GraphPad Prism 8 software and are depicted as mean ± SD. Significant differences between two groups were calculated using unpaired two-tailed t-test. A one-way AVOVA was performed to assess differences among multiple treatment groups unless mentioned otherwise. A two-way repeated measures ANOVA was performed to evaluate the effects of two factors (multiple treatment groups and time points). Dunnett’s or Games-Howell’s multiple comparisons test was applied in all ANOVA tests, besides qFISH analysis Sidak’s multiple comparisons test was used. Homogeneity of variances were confirmed used Brown-Forsythe test in one-way ANOVA and sphericity (Geisser-Greenhouse’s epsilon) in two-way repeated measures ANOVA. All data were tested positive for normality distribution using Kolmogorov-Smirnov test. Normality in qFISH data was assumed based on sample size of n≥84, following the Central Limit Theorem.

## Results

### ModRNA hTERT reactivates telomerase activity in MRC-5 cells

We first set out to produce modified hTERT mRNA (modRNA hTERT) that consists of CleanCap® Reagent AG as 5’ cap, 5’ and 3’ UTRs from the α-globin and β-globin genes, respectively, a segmented 120 bp poly (A) tail and the hTERT ORF (Fig. 1A,B). The modRNA consists of the modified nucleoside m1Ψ to reduce innate immune reaction and enhance stability and translational efficiency (Karikó, Buckstein, Ni, & Weissman, 2005; Nance & Meier, 2021). In parallel to hTERT, modRNA GFP was produced as control. Purified modRNAs for hTERT and GFP were loaded onto a denaturing gel to ensure the correct size of 3,708 bp for modRNA hTERT and 1034 bp for modRNA GFP (Supplement Fig. 1A). To validate the modRNA functionality we chose MRC-5, a fetal lung fibroblast cell line. 1 µg/mL modRNA GFP revealed a high transfection efficiency as GFP expression was detectable from 8 h to 120 h post-transfection (Supplement Fig. 1B). Since the GFP expression peaked between 24 h and 48 h these time points were chosen for further experimental analysis (Figure 1C). Next, we demonstrated a dose-dependent hTERT overexpression upon modRNA hTERT transfection (0.25 – 2 µg/mL) in the MRC-5 cells which possess negligible telomerase expression at baseline (Supplement Fig. 1C). To prove that the hTERT is translated into a catalytically active telomerase protein, telomerase repeat amplification protocol (TRAP) was performed. Strikingly, even the lowest concentration of modRNA hTERT 0.25 µg/mL was sufficient to induce robust telomerase activity in MRC-5 cells (Supplement Fig. 1D, E). Although the hTERT modRNA expression levels started to decrease at 48 h (Supplement Fig. 1C), the telomerase activity at 48 h was comparable to the 24 h time point.

**Figure 1.**
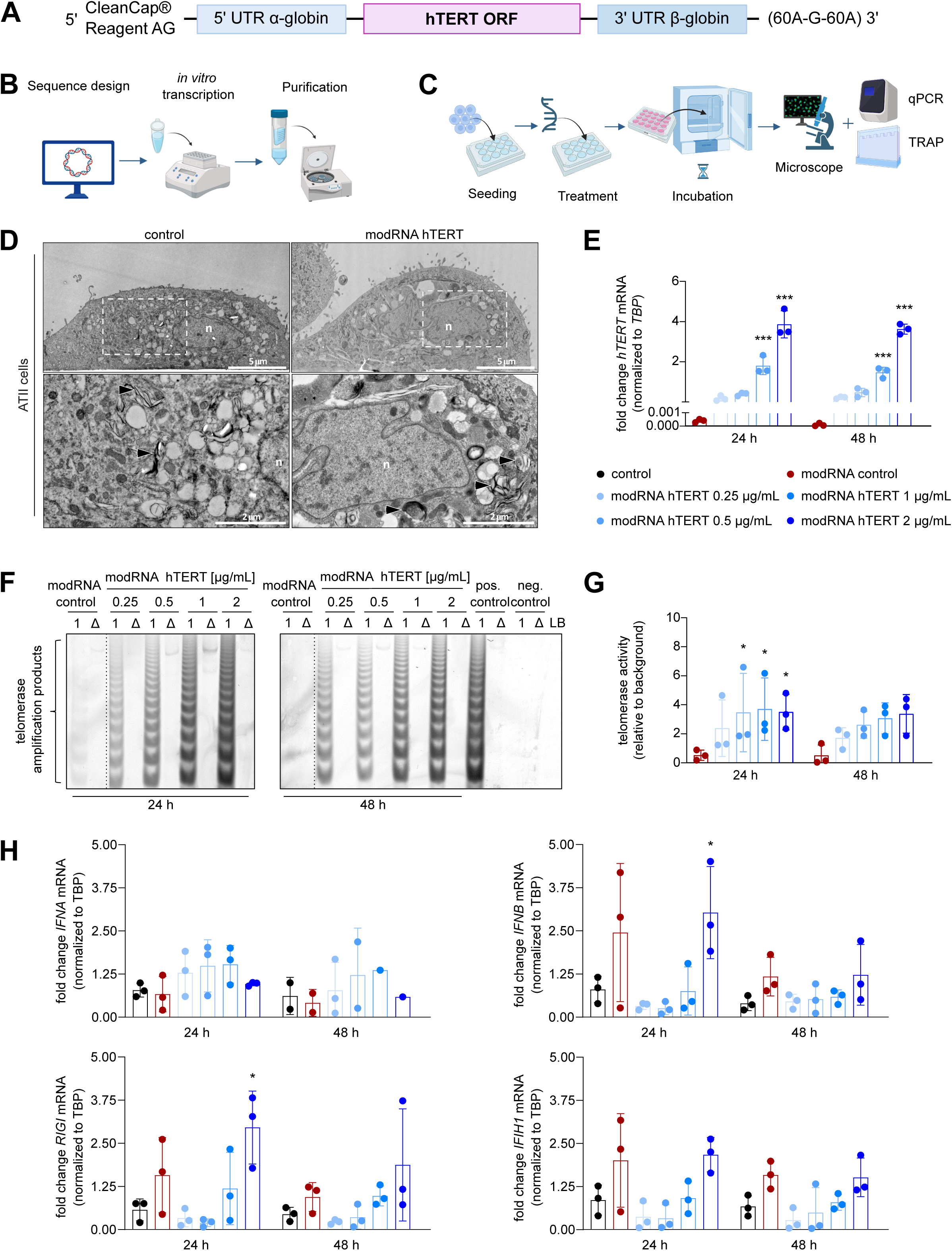
Dose-dependent increase of hTERT mRNA and telomerase activity *in vitro* after modRNA hTERT transfection. **A** Schematic presentation of the modRNA hTERT template. **B** The modRNA production is divided into three main steps: the sequence design, then *in vitro* transcription (IVT) where the modRNA is generated and lastly the purification step. **C** Schematic describing experimental setup used to validate transfection efficiency of modRNA GFP and functionality of modRNA hTERT. **D** Transmission electron microscopy of ATII cells. Arrowheads indicate lamellar body like structures. Nucleus is depicted as n. **E** Fold change of hTERT mRNA expression in ATII cells transfected with modRNA hTERT in a concentration range of 0.25 – 2 µg/mL compared to modRNA GFP 2 µg/mL control (modRNA control) after 24 h and 48 h (n = 3). **F** TRAP assay revealed reactivation of telomerase activity in ATII cells after 24 h and 48 h in dose-dependent effect. Positive (pos.) control = human induced pluripotent stem cells, negative (neg.) control = HUVEC cells, LB = lysis buffer, 1 = cell lysate, Δ = heat inactivated lysate. **G** Quantification of TRAP assays indicates a dose-dependent increase of telomerase activity compared to modRNA control (n = 3). **H** No prolonged immune reaction was detected after 48 h modRNA treatment in ATII cells compared to untreated control (n = 3). *p<0.05; ***p<0.001; Two-way ANOVA, Dunnett’s multiple comparisons test.

In summary, we successfully validated our optimized modRNA hTERT design and observed a dose-dependent overexpression on transcriptional and functional level in MRC-5. Thus, transient mRNA expression of modRNA hTERT, even at a minimal dosage is capable to induce telomerase activity.

### ModRNA hTERT treatment robustly induces telomerase activity in primary human alveolar type II pneumocytes

To validate and assess the therapeutic potential of modRNA hTERT in a more relevant model for lung disease, we utilized human primary ATII cells. First, we confirmed the cellular identity by assessing the ultrastructure through transmission electron microscopy highlighting the presence of lamellar bodies, specialized organelles that store and secrete surfactant. Notably, transfection of ATII cells with modRNA hTERT did not influence cell morphology (Figure 1D).

The efficiency of modRNA GFP transfection in ATII cells was consistent with those observed in MRC-5 cells (Supplement Fig. 2A). Moreover, transfecting ATII cells with the modRNA hTERT again showed a dose dependent increase in hTERT expression and telomerase activity (Figure 1E-G). Since it is well-established that “foreign” RNA can activate the innate immune system and induce an inflammatory response, we additionally evaluated the expression of several innate immune marker genes. Interferon α (IFNA) and interferon β (IFNB) are general markers for the activation of the innate immunity, while retinoic acid-inducible gene I (RIGI) and interferon induced with helicase C domain 1 (IFH1) are cytoplasmic sensors that detect foreign RNA and trigger the interferon signaling pathways. Only high dose of modRNA GFP (modRNA control) and modRNA hTERT (2 µg/mL) compared to untransfected cells (control) led to significant increase of IFNB and RIGI at 24 h, which quickly diminished at 48 h, indicating a transient and very brief immune reaction. IFNA and IFH1 expression remained unchanged (Fig. 1H). Taken together, we validated that modRNA hTERT treatment can significantly induce hTERT expression and reactivate telomerase activity in primary human ATII cells without provoking prolonged immune response.

### ModRNA hTERT enhances the proliferation capacity and improves cellular health

Next, we investigated whether the modRNA-mediated reactivation of hTERT expression and activity in ATII cells could lead to telomere elongation. To do so, we treated ATII cells with modRNA hTERT for 48 h and 96 h, followed by telomere qFISH analysis to quantify the telomere lengths. Telomere length at these time points did not increase, suggesting that the given times were too short or repetitive treatment would be necessary to achieve telomere elongation (Supplement Fig. 2B).

Therefore, modRNA hTERT was transfected in ATII cells at every second passage and we performed a quantitative growth curve analysis. This repetitive modRNA hTERT treatment resulted in a marked increase in population doubling level (PDL) by day 11, which continued to diverge significantly from the modRNA GFP treated control group until day 29 (fourth passage), indicating enhanced proliferative capacity after consecutive modRNA hTERT treatment (Fig. 2A; Supplement Fig. 2C). Increased PDL was accompanied by a significant decrease in mRNA levels of senescence markers such as cyclin-dependent kinase inhibitor 2A (CDKN2A), cyclin-dependent kinase inhibitor 1A (CDKN1A) and tumor protein (TP53) (Supplement Fig. 2D). To further substantiate these findings, we conducted a BrdU incorporation assay, which confirmed an increase in DNA synthesis, thereby validating the sustained proliferative activity driven by telomerase reactivation (Fig. 2B). In line, immunofluorescence analysis of cell proliferation markers Ki67 and phospho-histone H3 (pH3) showed a significant increase already after single treatment of modRNA hTERT (Fig. 2C). Conversely, γH2AX, a marker of DNA damage, was reduced after modRNA hTERT treatment. In contrast to the initial acute treatment, the long term and repetitive treatment led to a substantial elongation in telomere length, reinforcing the observed pro-survival effects (Fig. 2D, E). Collectively, our findings demonstrated that a double modRNA hTERT treatment is capable of enhancing proliferative capacity and reducing DNA damage, likely through telomere elongation, and ultimately improves the overall cellular fitness of ATII cells.

**Figure 2.**
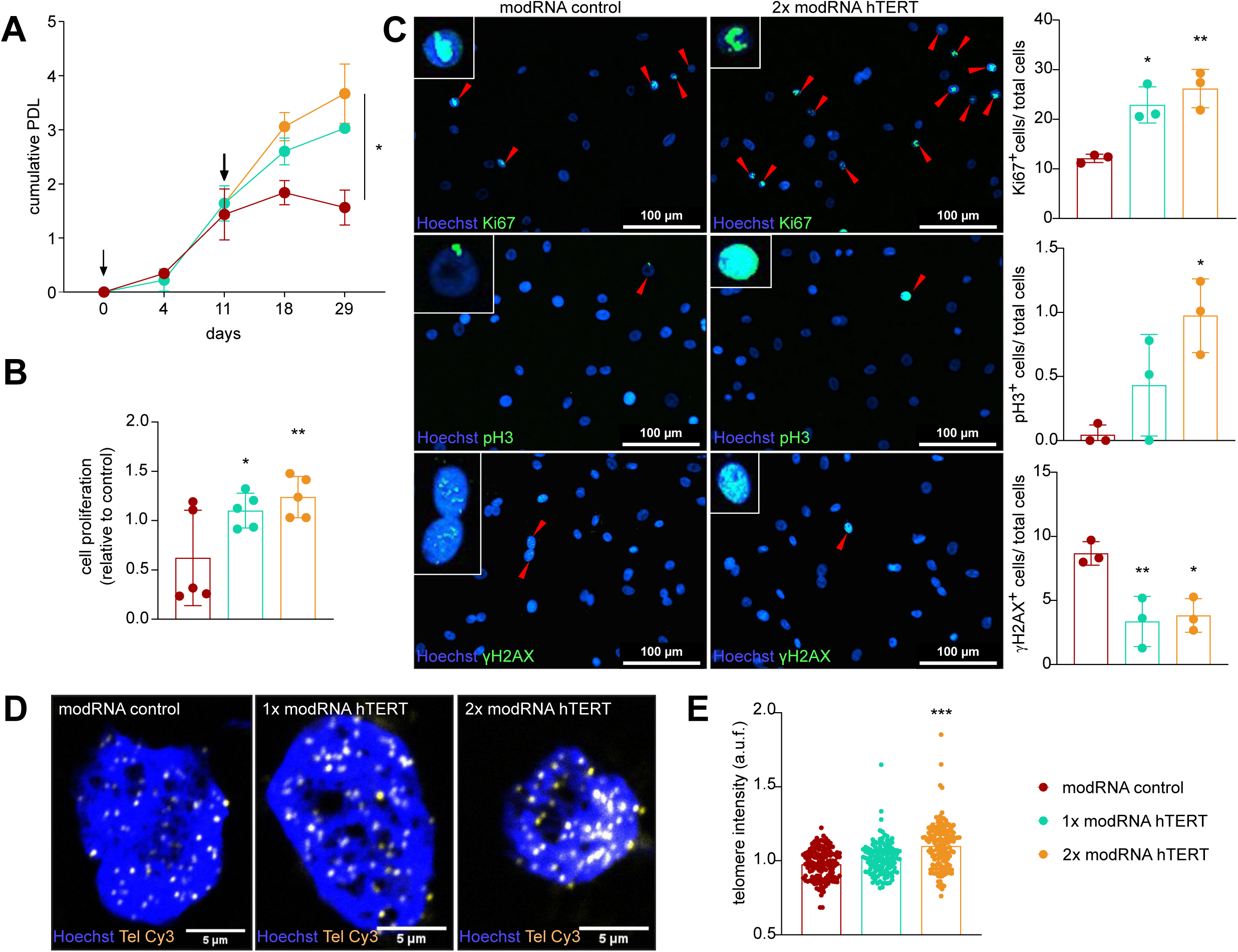
Repetitive modRNA hTERT treatment increased proliferation capacity and enhanced cell viability in ATII cells. **A** Population doubling level (PDL) demonstrated an increase after single modRNA hTERT treatment, which was further increased upon double modRNA hTERT treatment. Arrowheads indicate the time point of modRNA hTERT treatment performed in ATII cells (n = 3). Geisser-Greenhouse’s epsilon; Two-way ANOVA; Dunnett’s multiple comparison test. **B** BrdU assay indicated towards an increase in proliferation in ATII cells after modRNA hTERT treatment (n = 5). **C** Immunofluorescence staining of Ki67, pH3 and γH2AX, (green) revealed an increase in proliferation and a decrease in DNA damage after modRNA hTERT transfection in ATII cells (n = 3). **D** Representative images of qFISH staining in ATII cells after modRNA hTERT treatment. Brighter telomere spots (yellow, Tel Cy3), representing longer telomeres, were visible after 2x treatment with modRNA hTERT compared to 1x treatment and modRNA control group. Nucleus is depicted in blue (Hoechst). **E** Telomeric spots analysis revealed an increase of telomere length compared to modRNA control group (n ≥ 95 nuclei per group out of 3 biological replicates were imaged). Welch ANOVA; Games-Howell’s multiple comparisons test. *p<0.05; **p<0.01; ***p<0.001; One-way ANOVA or Two-way ANOVA, Dunnett’s multiple comparisons test.

### Self-spliced circular RNA hTERT as an alternative to prolong telomerase activity

We next sought to explore possibilities to extend the transient expression of modRNA hTERT by circularizing the *in vitro* transcription (IVT) product to increase stability thereby prolonging its therapeutic effects.

To produce circular RNA hTERT, the hTERT ORF was cloned into a vector adjacent to an internal ribosomal entry site (IRES). IRES-hTERT was then flanked by intron-exon sequences that serve as group I introns which induce circularization through self-splicing in the presence of the cofactors GTP and Mg^2+^ (Fig. 3A) (Wesselhoeft et al., 2018). In order to obtain a pure circular construct and to prove superior stability, IVT products were treated with exonuclease RNase R, which cleaved linear RNAs thereby enriching circular RNAs (Supplement Fig. 3B). Reverse transcription and PCR amplification with divergent primers was performed to validate the circular RNA hTERT. The PCR amplicon was sequenced to confirm the expected splice site and circularized sequence (Fig. 3B; Supplement Fig. 3B). Circular RNA GFP (circular RNA control) was transfected into HEK293 cells to assess transfection efficiency and protein translation potential (Supplement Fig. 3C). After validating successful transfection and GFP expression in HEK293, the modRNA hTERT (linear RNA) was transfected in parallel with circular RNA hTERT in MRC-5 cells. At 24 h post-transfection, the levels of linear and circular hTERT transcripts were comparable. However, 48 h post transfection the level of linear RNA hTERT transcript started to decline, while the circular RNA transcript levels remained stable (Fig. 3C). These observations were reinforced by TRAP assay (Fig. 3D) where telomerase activity at 24 h was similar, but at 48 h the circular RNA hTERT treated group presented a significantly higher telomerase activity compared to the linear RNA hTERT (Fig. 3E). Strikingly, in contrast to modRNA hTERT, circular RNA hTERT, resulted in significant telomere elongation after 96 hours of treatment (Fig. 3F-G). To conclude, we provide pioneering results for the testing and validation of circularized hTERT RNA as an alternative to linear modRNA hTERT constructs to reactivate telomerase expression and activity.

**Figure 3.**
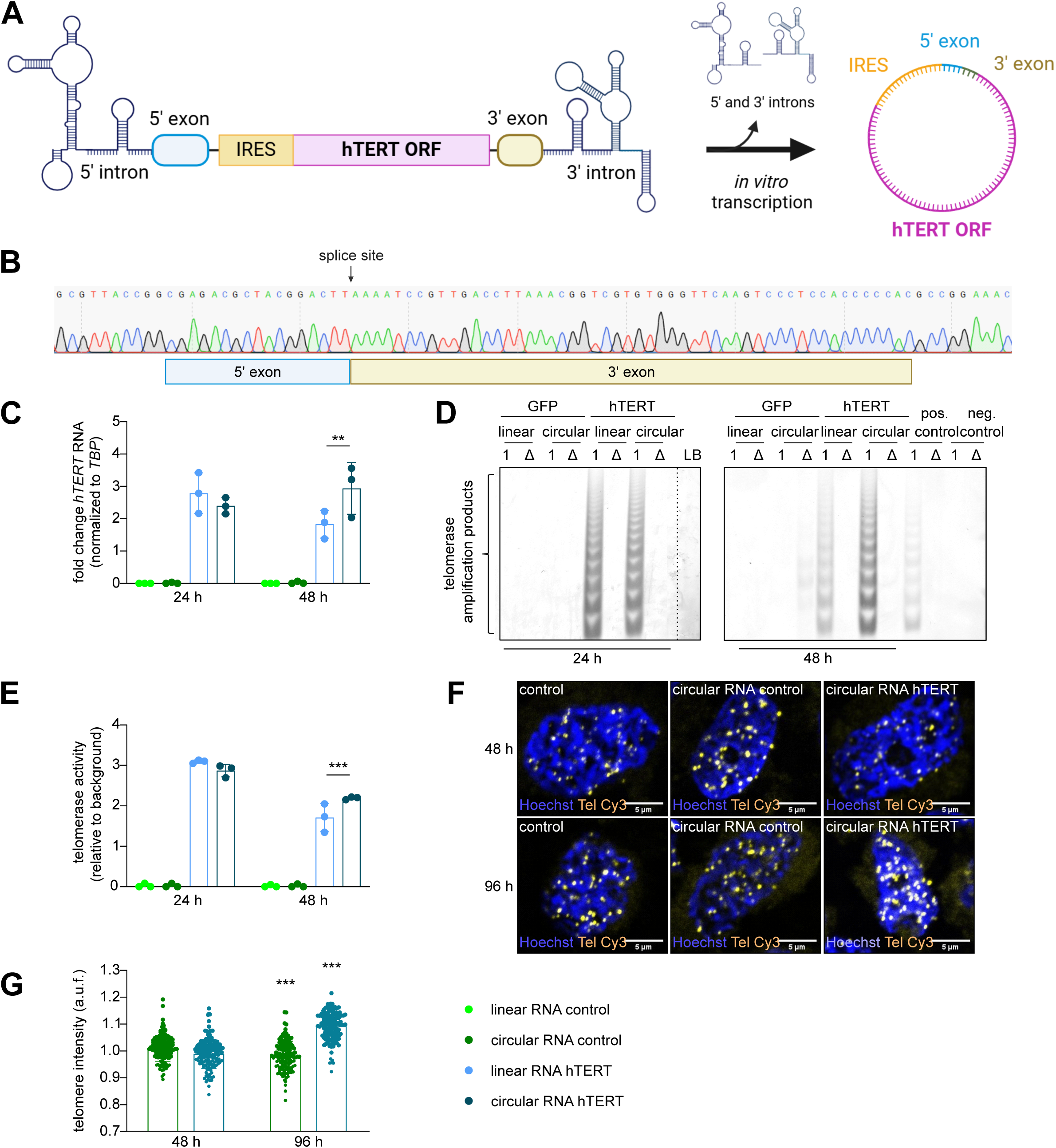
Circular RNA hTERT is more stable, resulting in a prolonged telomerase activity compared to linear RNA hTERT. **A** Schematic presentation of the circular RNA hTERT construct design. **B** Sanger sequencing of the hTERT PCR product, obtained after reverse transcription of circular RNA using divergent primers, revealed the circularization of the exons with expected splice site. **C** Transcript levels of hTERT comparing linear and circular RNA hTERT treatments after 24 h and 48 h in MRC-5 cells. (n = 3). **D** TRAP assay confirmed the telomerase activity level after transfection of both linear and circular hTERT RNA in MRC-5 cells. Positive (pos.) control = HEK293 cells, negative (neg.) control = HUVEC cells, LB = lysis buffer, 1 = lysate, Δ = heat inactivated lysate. **E** Quantification of TRAP assays indicating an increase of telomerase activity after circular RNA in comparison to linear RNA at 48 h timepoint (n = 3). **F** Representative images of qFISH staining in MRC-5 cells after circular RNA hTERT transfection. Brighter telomere spots (yellow, Tel Cy3), representing longer telomeres, were visible after 96 h treatment with circular RNA hTERT. Nucleus is depicted in blue (Hoechst). **G** Analysis of telomeric spots revealed an increase of telomere length in circular RNA hTERT treatment group at 96 h compared to circular RNA control group (n ≥ 132 nuclei per group out of 3 biological replicates were imaged). **p<0.01; ***p<0.001; One-way ANOVA or Two-way ANOVA, Dunnett’s multiple comparisons test.

### ModRNA hTERT reduces senescence in human precision-cut lung slices derived from patients with idiopathic pulmonary fibrosis

Following up on the pro-proliferative and pro-survival effects conferred by the modRNA hTERT treatment on cells *in vitro* we aimed to test its potential in a state-of-the-art, patho-physiologically and clinically relevant platform. We employed human PCLS derived from explanted lungs, which exhibited end-stage PF (Table 1, Fig. 4A). First, we assessed the expression timeline after modRNA hTERT transfection from a non-fibrotic patient (Table 1) and two PF patients (Table 1), where a strong induction of hTERT mRNA was observed after 24 h (Fig. 4B) and 48 h (Supplement Fig. 4A). To evaluate the efficacy of modRNA hTERT therapy, PCLS from four PF patients diagnosed with either secondary fibrosis or IPF were employed. Notably, LDH levels released between the modRNA control and modRNA hTERT treatment did not show any statistical difference, indicating optimal viability throughout the entire experimental period of 96 h (Fig. 4C). Presence of senescent alveolar epithelial cells in PF patients contributes to impaired regeneration, chronic inflammation, and fibrosis (Parimon et al., 2023). Strikingly, levels of senescence-associated β-galactosidase (SA-β-gal) was significantly lower after modRNA hTERT compared to modRNA control which exhibited strong SA-β-gal staining at 96 h after transfection (Fig. 4D, E). Subsequently, supernatants were collected to evaluate SASP-related inflammatory responses and to analyze fibrosis markers in these PF-derived PCLS after modRNA hTERT treatment. Notably, 96 h after modRNA hTERT treatment the inflammatory response of IL6 and IL8, as well as the fibrotic TGFβ signaling and the fibrosis marker pro-COL1A1 were significantly reduced (Fig. 4F).

**Figure 4.**
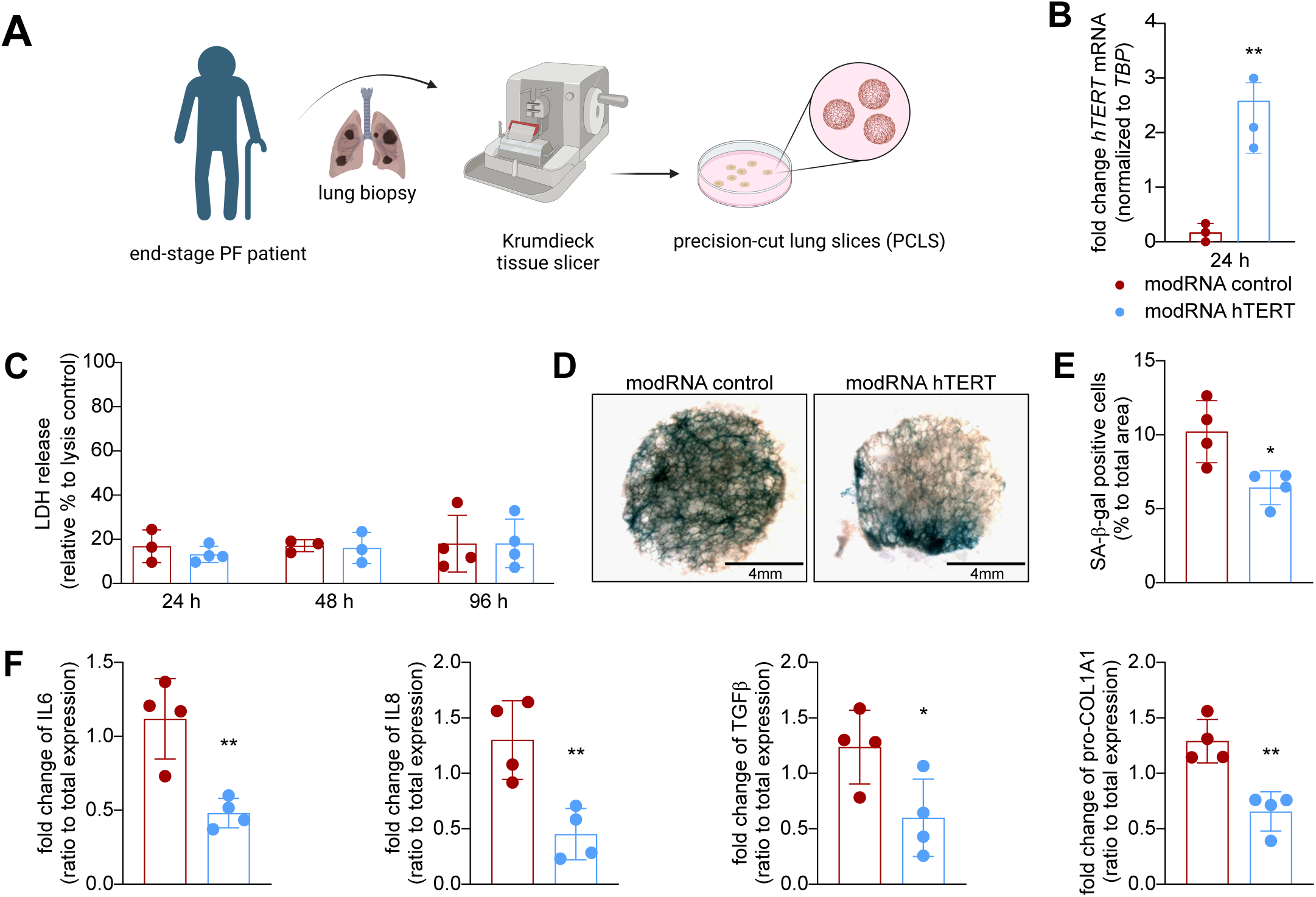
Therapeutic modRNA hTERT treatment in PF patient-derived PCLS leads to reduced senescence, inflammation and fibrosis. **A** Schematic presentation of PCLS (precision-cut lung slices) preparation. PF: pulmonary fibrosis; **B** Validation of overexpression of hTERT transcript level at 24 h after modRNA hTERT treatment in PCLS (n = 4). **C** LDH release is given as % of triton-lysed control (set to 100 %). No increase in LDH release was detected after modRNA hTERT compared to modRNA control (n = 3-4). **D** Representative images of SA-β-gal stained PCLS (n = 4). SA-β-gal positive region in the PCLS tissues are in blue color. **E** Quantification of area occupied by SA-β-gal positive staining within the PCLS (N = 4). **F** Downregulation of the IL6, IL8, TGFβ and pro-COL1A1 protein level relative to total expression after modRNA hTERT treatment (n = 4). *p<0.05; **p<0.01; unpaired t test; Two-way ANOVA, Dunnett’s multiple comparisons test.

Taken together, these findings confirmed that modRNA hTERT treatment in human end-stage PF-PCLS conferred anti-senescence effects without inducing cytotoxicity. Our data suggest general therapeutic potential of modRNA hTERT against fibrotic interstitial lung diseases, including but not limited to IPF, as it effectively reduced inflammatory and fibrotic responses.

## Discussion

We report a novel treatment option for PF by employing modRNA technology to transiently reactivate telomerase activity in ATII cells. Accounting for the most aggressive form of ILDs, IPF occurs with the highest frequency in ILD patients (Maher, 2024; Rea et al., 2024). One risk factor for IPF is the premature shortening of telomeres in the lung (Van Batenburg et al., 2020). Povedano et al. already demonstrated in rodents that AAV9-Tert could be utilized as a therapeutic option to ameliorate PF induced by bleomycin treatment in mice with short telomeres (Povedano et al., 2018). Of note, bleomycin induced fibrosis in mice is not progressive and therefore this model does not recapitulate the full complexity of PF. Liu et al. performed a conditional knockdown of TERT in ATII cells combined with a bleomycin stress treatment. Although they observed signs of lung fibrosis, there were no significant effects on telomere lengths in these mice (Liu et al., 2019). This might have occurred since laboratory bred mice have extremely long telomeres and recent efforts have been made to generate mice with telomeres comparable to humans, highlighting that most studies in *in vivo* mice models have limited translatability (Calado & Dumitriu, 2013; Smoom et al., 2023). Moreover, AAV gene therapy strategies have some limitations. Firstly, the existence of neutralizing antibodies against naturally occurring AAV in up to 70% of the population, represents a major clinical barrier. (Wang, Gessler, Zhan, Gallagher, & Gao, 2024). Secondly, due to the constitutive and long-term overexpression provided by AAVs, its use for overexpressing hTERT in patients might lead to regulatory concerns due to the potential risk for tumorigenesis. By contrast, due to their non-integrative and transient expression profile, modRNAs appear to be an ideal vector for hTERT delivery in patients (Sahin, Karikó, & Türeci, 2014). Additionally, the modRNA production using IVT method is scalable and cost efficient thus making it an affordable therapy compared to AAV gene therapy (Chaudhary, Weissman, & Whitehead, 2021). One groundbreaking example of this modRNA technology is the COVID-19 vaccine development which highlights the clinical readiness of this drug class (Warne et al., 2023).

In our study, both *in vitro* (MRC-5 cells as wells as primary ATII cells) and *ex vivo* human end-stage PF-PCLS tissue were successfully transfected with modRNA hTERT leading to a transient expression of the functionally active telomerase protein. The catalytically active protein increased the proliferative capacity, decreased DNA damage while significantly increasing the telomere length in the treated ATII cells. This observation is in line with a previous study by Ramunas et al., where they applied modRNA hTERT *in vitro* in a non-disease setting to increase the proliferation capacity in lung fibroblasts and primary skeletal muscle myoblasts (Ramunas et al., 2015). It is important to acknowledge that these general benefits could be the result of other telomerase-related function aside from telomerase-induced telomere elongation. Apart from its canonical role, hTERT it is also known to possess non-telomeric functions in the mitochondria (Chatterjee et al., 2021). Thus, there might be other non-canonical functions of hTERT due to which primary lung cells can benefit after hTERT treatment. This aspect should be further investigated in the context of transient modRNA hTERT therapy for lung fibrosis. To overcome the various limitations of available mouse models and *in vitro* cell lines for investigation of PF, we utilized PCLS, a state-of-the-art organotypic model of the human lung. Particularly, we used viable diseased lung tissue from patients suffering from end-stage PF. These PCLS retain the complex pathogenesis of PF by preserving active cellular and mechanical function of various cell types in their native lung architecture (Hesse et al., 2022; Koziol-White, Gebski, Cao, & Panettieri, 2024; Neuhaus et al., 2018, 2017). This model enables *ex vivo* investigation of integrated cellular response to various treatment options, including modRNA hTERT, to support pre-clinical tests and toxicity studies. Our data reveals clear downregulation of SA-β-gal activity within the tissue as well as SASP-related inflammatory responses highlight the therapeutic potential of modRNA hTERT in mitigating inflammation and fibrosis in PF.

It is commonly observed that isolated ATII cells undergo spontaneous differentiation towards ATI cells, which leads to a reduction in their proliferative capacity (Strunz et al., 2020). In elderly patients, the lung ATII cells also exhibit advanced stages of senescence in lung fibrosis (Parimon et al., 2023). In this context we showed that modRNA hTERT treatment has the ability to rescue ATII cells from senescent state, since we observed the modest reduction of the senescence markers CDKN2A, CDKN1A and TP53 after modRNA hTERT treatment at later passages *in vitro* and a significant reduction in β-gal staining in human PCLS tissue *ex vivo*. In line with this finding, we could demonstrate that modRNA hTERT treatment was able to alleviate the secretion of SASP pro-inflammatory markers IL6 and IL8 along with reduced levels of TGFβ and pro-COL1A1, which are key mediators for fibrosis progression. This highlights the potential of modRNA hTERT to prevent senescence, leading to a decrease of inflammation as well as mediators of fibrosis.

The human innate immune system has several pattern recognition receptors that detect foreign RNA and activate downstream signaling cascades to trigger an immune reaction, like TLR7, RIGI or IFH1(Fang et al., 2022; Nance & Meier, 2021). We demonstrated that after treatment with modRNA hTERT *in vitro*, there were no prolonged upregulation of immune markers, such as RIGI, IFH1, IFNA and IFNB. These results are in alignment with the immune safety analysis of the FDA-approved modRNA vaccine (BNT162b2) against COVID-19, where systemic reaction due to the vaccine were resolved within one to two days (Polack et al., 2020). Finally, in an attempt to prolong the transient expression of hTERT, we produced a circular, exonuclease-resistant form of RNA, which provided higher stability and longer window of modRNA hTERT therapeutic efficacy as evidenced by a significant telomere elongation. To our knowledge this is the first study that uses circularized IVT hTERT for successful telomerase reactivation. In summary, our findings support the therapeutic potential of repeated modRNA-mediated hTERT therapy or circular RNA hTERT therapy for PF treatment by transient yet significant elevation of telomerase expression and activity which is sufficient to elongate telomeres, and reduce senescence along with increased cellular health in primary lung cells and organotypic tissue from end-stage PF, including IPF.

## Acknowledgements

We thank A. Gietz, I. Riedel, N. Lehmann and E. Westerheide from the Institute of Molecular and Translational Therapeutic Strategies (IMTTS), Hannover Medical School (MHH) for excellent technical support. We appreciate the support from the Central Laser Microscopy Unit, MHH. Additionally, we thank Taihua Yang, PhD lab from Department of Liver Surgery, Renji Hospital, School of Medicine, Shanghai Jiao Tong University in China for kindly providing plasmid for circular RNA generation used in this study. Some schematic figures were created with Biorender.com.

## Author Contributions

JY designed and performed the experiments with subsequent analysis. KG and JY performed the human lung tissues experiments. DL, MJ, ST and LO contributed to the modified RNA and circular RNA methods. CBr performed electron microscopy experiments. MJo contributed to RNA isolation from PCLS. KG, PZ, CW, CH and KS provided human lung tissues. KG and CH performed LDH and ELISA assays in PCLS. CB conceived the original idea of the study and supported manuscript preparation. TT, KS, ST, CBr, SC and CB received funding. SC was involved in training, supervision and experimental design of this project. JY, CB and SC wrote the manuscript. CBr, CH, KS, ST, TT, and CB revised the manuscript. All authors reviewed and approved the final version of the manuscript.

## Supplementary Figures

**Supplementary Figure 1.**
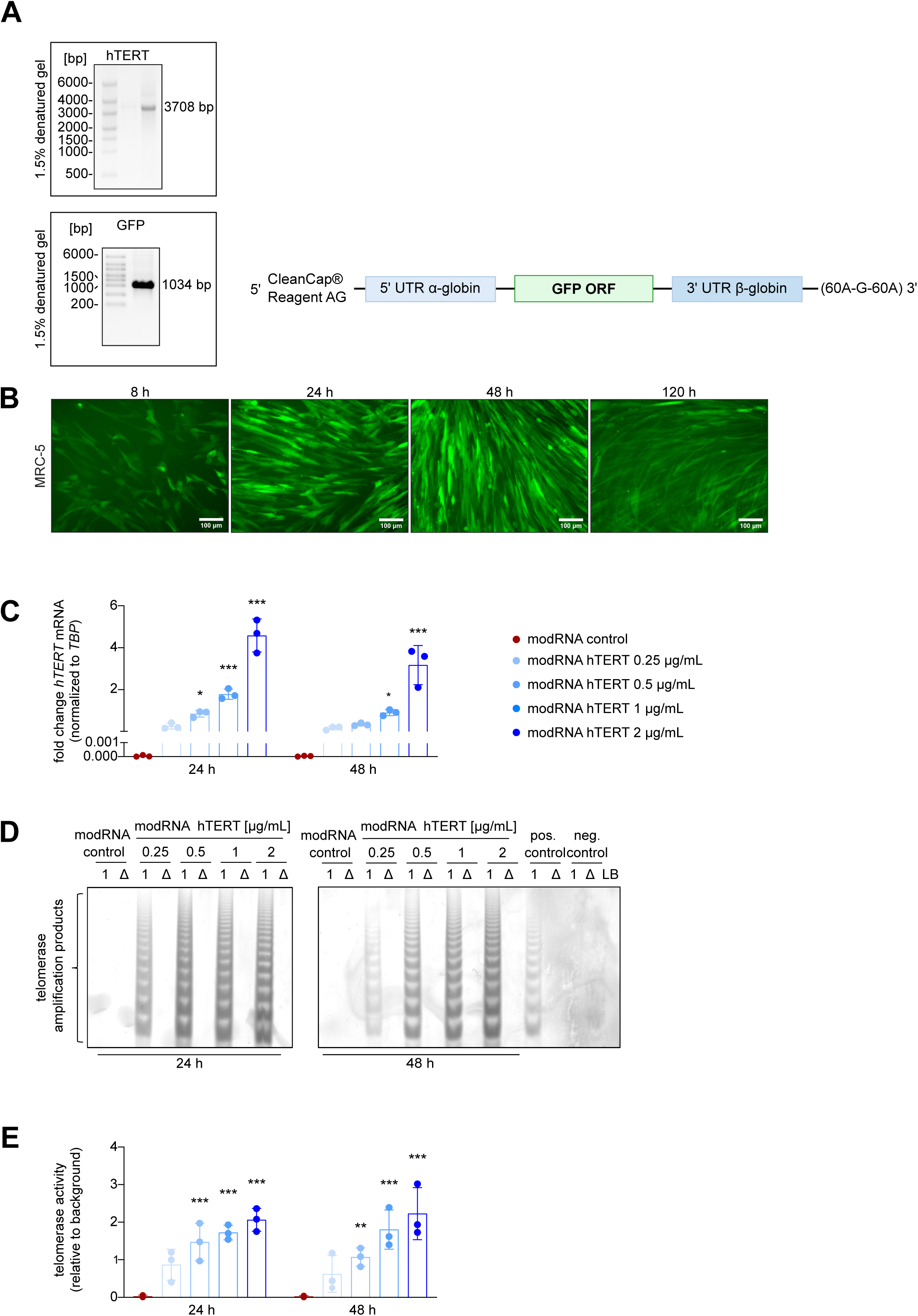
Dose-dependent increase of hTERT mRNA and telomerase activity *in vitro* after modRNA hTERT transfection in MRC-5 cells. **A** Validation of the hTERT and GFP modRNA product size after IVT in 1.5 % denatured gel. ModRNA GFP is generated by substituting hTERT ORF with the GFP coding sequence. **B** Images showing the successful transfection of modRNA GFP 2 µg/mL and stable GFP expression for until 48 h and a significant decline at 120 h in MRC-5 cells. **C** Fold change of hTERT mRNA expression after transfection with modRNA hTERT in a concentration range of 0.25 – 2 µg/mL compared to modRNA control (modRNA GFP 2 µg/mL) after 24 h and 48 h in MRC-5 cells (n = 3). **D** TRAP assay revealed telomerase activity after 24 h modRNA hTERT transfection in MRC-5 cells. Positive (pos.) control = HEK293 cells, negative (neg.) control = HUVEC cells, LB = lysis buffer, 1 = lysate, Δ = heat inactivated lysate. **E** Quantification of TRAP assays in MRC-5 (n=3). *p<0.05; ***p<0.001; Two-way ANOVA, Dunnett’s multiple comparisons test.

**Supplementary Figure 2.**
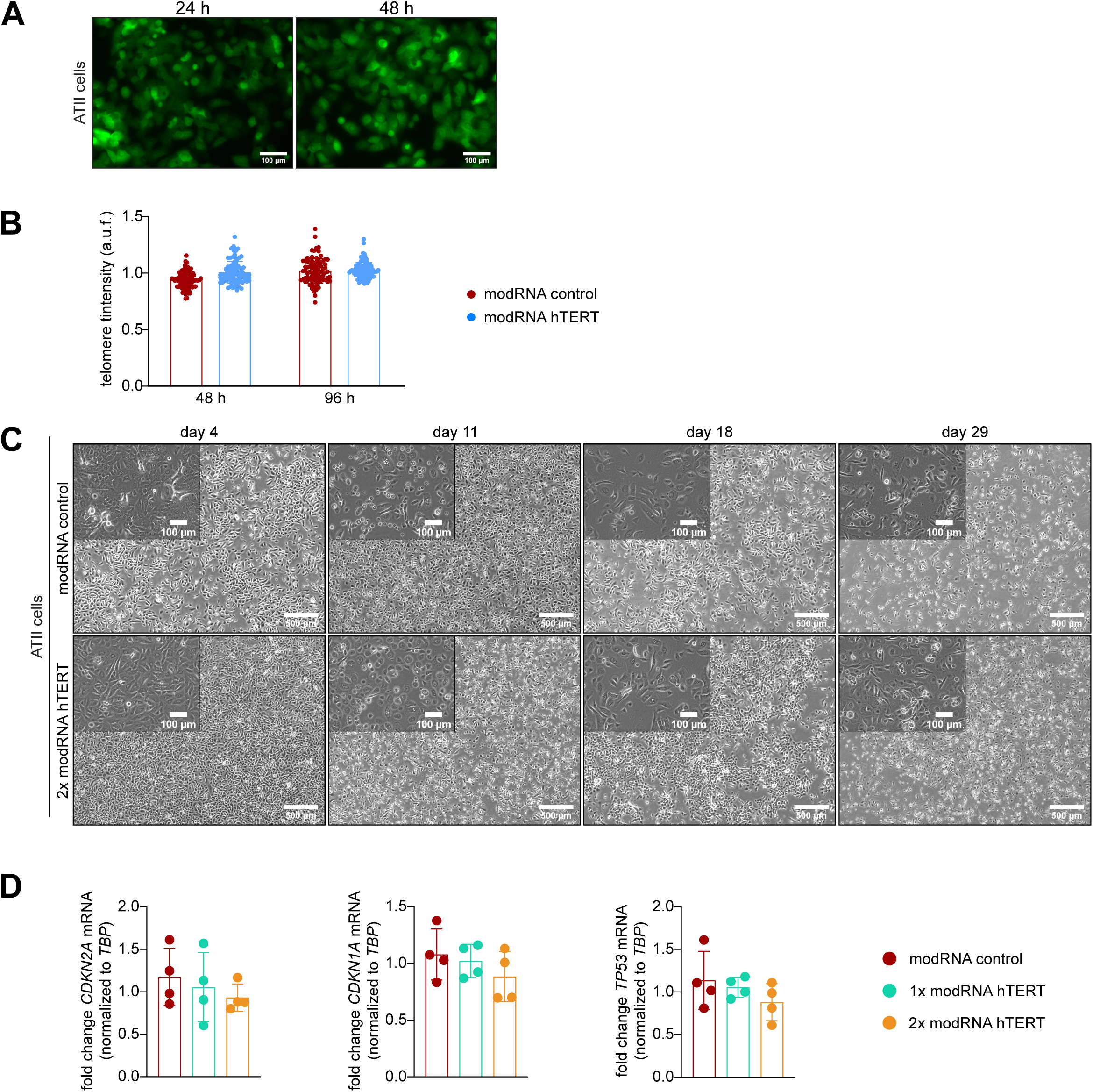
Repetitive treatment of modRNA hTERT increase proliferation capacity and decrease expression of senescence-related marker in ATII cells. **A** Images showing successful transfection of modRNA GFP 2 µg/mL in ATII cells as observed by GFP expression in 24 h an 48 h post transfection. **B** Telomere qFISH analysis demonstrated no telomere elongation after 48 h and 96 h modRNA hTERT treatment in ATII cells (n ≥ 84 nuclei per group out of 3 biological replicates were imaged) modRNA GFP is used as the modRNA control. **C** Representative brightfield images on the day of passaging in ATII cells of modRNA control compared to 2x treatment modRNA hTERT. **D** A decreasing trend of senescence-related markers (CDKN2A, CDKN1A, TP53) was detected after two doses of modRNA hTERT treatment (n=4). *p<0.05; One-way ANOVA or Two-way ANOVA, Dunnett’s or Sidak’s multiple comparisons test.

**Supplementary Figure 3.**
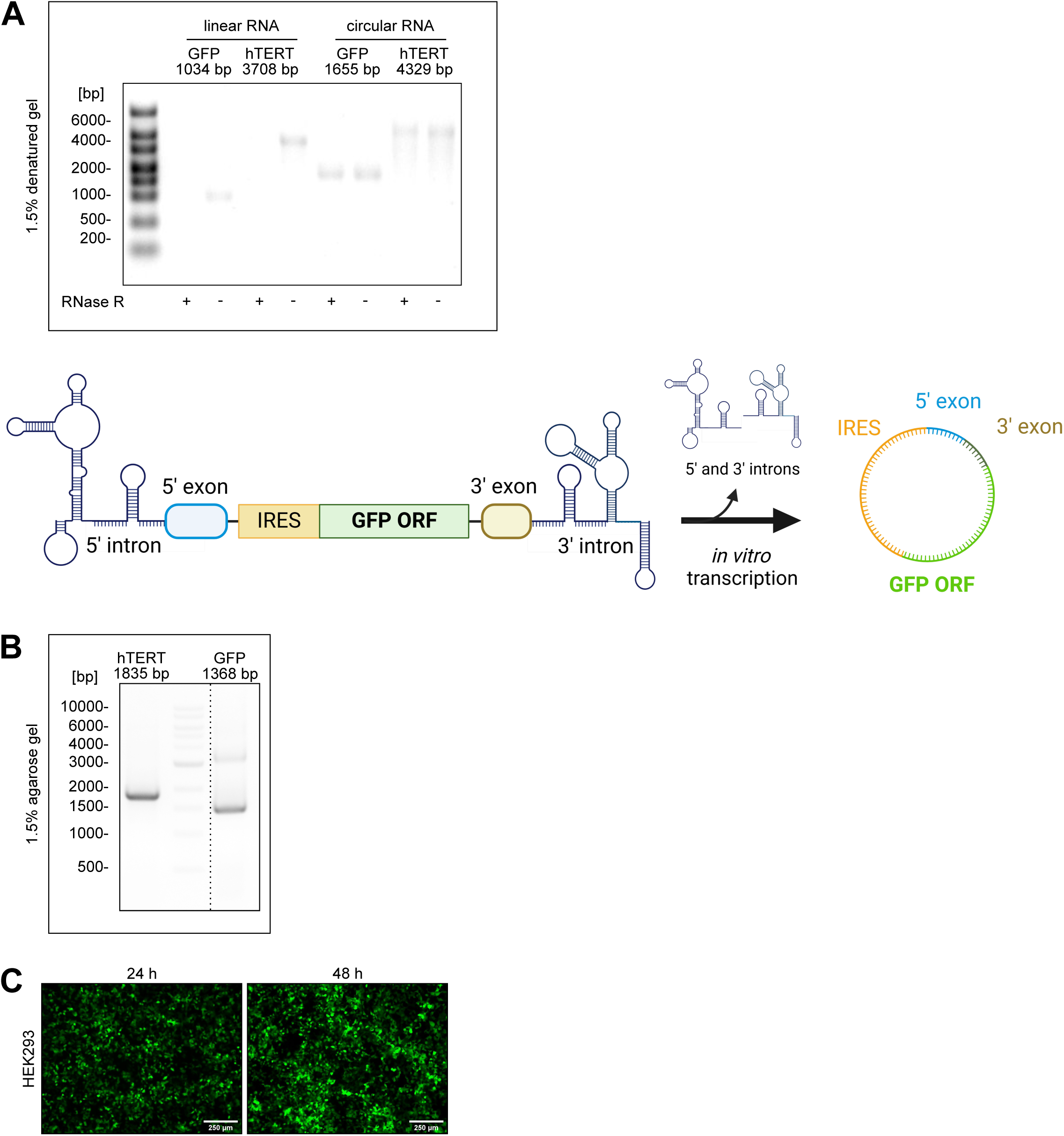
Validation of circular RNA. **A** Validation of the size of produced linear and circular RNA products using IVT encoding for hTERT or GFP. To prove superior stability of circular RNA the linear and circular RNAs were treated with RNase R and loaded on 1.5 % denatured gel. Circular GFP is generated by substituting hTERT ORF with the GFP coding sequence. **B** Validation of circularization by performing PCR with divergent Primer of reverse transcribed circular RNA. Loaded on 1.5 % agarose gel. **C** Representative images confirming successful transfection of circular RNA GFP 2 µg/mL in HEK293 cells.

**Supplementary Figure 4.**
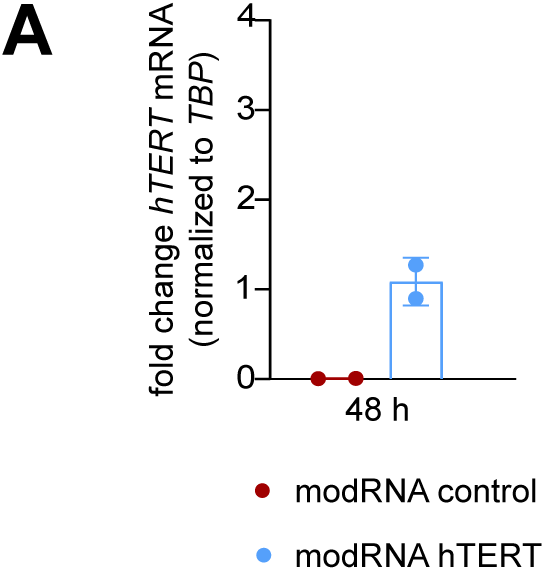
Overexpression of hTERT mRNA after modRNA hTERT transfection in PCLS. **A** Fold change of hTERT mRNA expression after 48 h of transfection with modRNA hTERT (1 µg/mL) compared to modRNA GFP (modRNA control) (1 µg/mL) in PCLS (n = 2).

